# Mitochondrial cytochrome c accumulation accompanies reduced electron flux through complex IV without enhancing cell sensitivity to apoptosis

**DOI:** 10.64898/2026.06.26.733950

**Authors:** Alexander V. Zhdanov, Nadezda A. Brazhe, Evelina Nikelshparg, Luke Power, Philip Lewis, Paula V. Silva, Marcel van de Wouw, Patrick B. F. O’Connor, John F. Cryan, Olga V. Sosnovtseva, Dmitry E. Andreev, Martina M. Yordanova, Pavel V. Baranov, Ruslan I. Dmitriev, Dmitri B. Papkovsky

**Author notes:** Corresponding author: Dr. Alexander V. Zhdanov, Pharmacy Building, College Road, Cork, Ireland.

## Abstract

We show that chronic impairment of mitochondrial respiration is associated with marked accumulation of cytochrome c (Cyt*c*) protein. Using SCO2-deficient HCT116 cells lacking functional cytochrome c oxidase and wild-type cells exposed to sustained hypoxia, we found that substantial mitochondrial Cyt*c* accumulation parallels reduced electron flux through Cyt*c*. SCO2-deficient cells exhibited equally elevated Cyt*c* levels under normoxia (19% O_2_) and hypoxia (0.1-3% O_2_). Wild-type cells under sustained hypoxia accumulated Cyt*c*, reaching levels comparable to those in SCO2-deficient cells. This effect was reversible upon reoxygenation. Increased Cyt*c* protein levels were also observed in other cell models, including primary cortical neurons cultured under chronic hypoxia and in cerebral cortex tissue from hypoxia-exposed mice. Cyt*c* accumulation occurred independently of *CYCS* transcription, mRNA translation, HIF activation, ROS production and changes in mitochondrial network. Pharmacological inhibition of complex III was likewise accompanied by increased Cyt*c* levels, whereas mitochondrial uncoupling had no effect, suggesting that impaired electron transfer rather than membrane depolarisation per se underlies this association. Raman spectroscopy revealed enrichment of reduced Cyt*c* and an increased Cyt*c*-to-cytochrome b ratio in respiration-deficient cells. Further supporting a stabilisation-based mechanism, the fraction of membrane-unbound ferro-Cyt*c* was decreased in SCO2-deficient cells, consistent with moderate cardiolipin enrichment, which is known to enhance retention of Cyt*c* at the inner mitochondrial membrane. Despite elevated mitochondrial Cyt*c* content, SCO2-deficient cells were less susceptible to apoptosis induced by intermittent hypoxia or dichloroacetate. Together, these findings indicate that reduced electron flux through complex IV is associated with Cyt*c* accumulation through increased protein stability and membrane retention without enhancing apoptotic sensitivity.

## Introduction

Cytochrome c (Cyt*c*) is a globular 12 kDa heme-protein involved in many fundamental processes, including transport of electrons and OxPhos, free radicals turnover, maintenance of cellular redox balance, apoptosis and intercellular signalling [1–4]. Although under certain physiological and pathological conditions Cyt*c* can function as an extracellular signalling molecule [2], it is predominantly localised within the mitochondrial intermembrane space, where it loosely associates with the inner mitochondrial membrane through electrostatic interactions. Within the electron transport chain (ETC), Cyt*c* undergoes continuous cycles of reduction and oxidation while transferring electrons from coenzyme Q : cytochrome c reductase (complex III) to COX (complex IV). In addition to its role as a mobile electron carrier, oxidised ferri-Cyt*c* can accept electrons from superoxide radicals and hydrogen peroxide, redirecting them back into the ETC and thereby contributing to ROS detoxification and redox balance [5]. Furthermore, upon binding to the mitochondrial phospholipid CL, ferri-Cyt*c* acquires peroxidase activity, catalysing CL oxidation—an event recognised as critical for mitochondrial membrane remodelling and the initiation of apoptosis [6].

Electron transfer from Cyt*c* to molecular oxygen via COX is generally considered a rate-limiting step of the ETC under physiological conditions. COX activity is tightly regulated by multiple factors, including oxygen availability, ΔΨm polarisation, intramitochondrial Ca²⁺ levels, and the cellular ATP/ADP ratio [7]. Notably, COX exhibits an exceptionally high affinity for oxygen, such that oxygen concentration becomes limiting for electron transfer through Cyt*c* only under conditions of severe hypoxia [8].

Encoded by nuclear genomic DNA, apo-Cyt*c* is synthesised in the cytoplasm and subsequently translocated across the outer mitochondrial membrane into the intermembrane space. There, apo-Cyt*c* is converted into its mature holo-form through the covalent attachment of a heme prosthetic group. This maturation step strictly requires reduction of the two conserved cysteine residues in apo-Cyt*c* and the ferrous state of the heme iron, enabling their utilisation as substrates by holo-Cyt*c* synthase [9].

Beyond heme incorporation, Cyt*c* undergoes several post-translational modifications (PTMs) that regulate its conformation, activity, and cellular fate. Phosphorylation at Thr^28^, Ser^47^, Tyr^48^, and Tyr^97^ attenuates Cyt*c* function both as an electron carrier and as an inducer of apoptosis, with phosphorylation at Tyr^48^ serving as a prominent example [10]. Free radical–dependent nitration of tyrosine residues (Tyr^67^, Tyr^74^, and Tyr^97^) enhances Cyt*c* peroxidase activity while suppressing its pro-apoptotic function [9]. Acetylation further modulates the redox properties of Cyt*c*, affecting its propensity to undergo reduction and oxidation cycles Collectively, these PTMs influence Cyt*c* structure, stability, and affinity for interaction partners, particularly CL. In its CL-bound state, Cyt*c* is largely excluded from electron transfer within the ETC [11]. An increased fraction of CL-bound Cyt*c* has recently been reported in several tumour types, in which OxPhos could be suppressed in a Cyt*c*–CL–dependent manner [12].

The redox state of Cyt*c* itself is a key determinant of its three-dimensional structure, interaction network, and potentially its intracellular turnover, as well as its role in apoptosis following release into the cytosol. While the ability of reduced Cyt c to bind Apaf-1 and promote apoptosome assembly remains debated, the oxidised form of Cyt c is generally regarded as a potent trigger of apoptosis. [13–15]. In addition, apo-Cyt*c* has been reported to inhibit apoptosome formation by competing with holo-Cyt*c* for Apaf-1 binding [16].

Although data on Cyt*c* degradation rates are limited, the protein is generally considered as relatively stable. Its turnover is tissue-specific and has been reported to decrease following release from mitochondria into the cytosol. In muscle tissue—the best characterised model—the half-life of Cyt*c* ranges from 4 to 10 days [17]. Expression of the Cytc-encoding gene *CYCS* and intracellular Cyt*c* levels are tissue-specific, with reported concentrations on the order of ∼0.1 nmol per mg of cellular protein in rat cardiomyocytes [18]. While Cyt*c* abundance is typically tightly regulated, mitochondrial Cyt*c* levels can change under conditions of altered metabolic demand. For instance, activation of the *CYCS* proximal enhancer by CREB and NRF1 has been shown to increase Cyt*c* expression in response to elevated neuronal activity in hippocampal neurons in vitro [19]. Despite extensive investigation of Cyt*c* function, the effects of oxygen availability, energy demand, and metabolic stress on Cyt*c* dynamics remain poorly understood. Notably, chronic hypoxia in pregnant rats was found to increase Cyt*c* levels in foetal heart by approximately 70%, while reducing Cyt*c* levels in foetal liver by 54% [20].

Previously, we reported increased Cyt*c* protein levels in the culture of pheochromocytoma PC12 and neuroblastoma SH-SY5Y cells, adapted to one-month chronic ‘physiological’ hypoxia [21]. This data suggests that Cyt*c* abundance may be dynamically regulated under conditions that restrict electron transfer through the ETC.

However, despite extensive knowledge of Cyt*c* structure and function, it remains unclear whether Cyt*c* accumulation represents a general adaptive response to limited O_2_ availability, ETC dysfunction or metabolic state, and how it influences cellular responses to proapoptotic stimuli.

To address these questions, we investigated Cyt*c* regulation in several cellular and animal models characterised by limited O_2_ availability and / or reduced electron transfer capacity. These included, alone with hypoxic animal tissue, primary cortical neurons and HCT116 cells with intact and pharmacologically inhibited mitochondrial function, also HCT116 cells lacking the SCO_2_ gene (synthesis of cytochrome C oxidase 2), which are unable to assemble functional COX [22, 23]. By combining these models, we sought to gain new insight into the adaptive and pathophysiological roles of Cytc under conditions of mitochondrial stress. Specifically, we examined how O₂ levels, respiratory chain competence, and metabolic demand shape Cytc abundance, and whether alterations in Cytc content influence cellular responses to proapoptotic stimuli.

## Materials and Methods

### Reagents

MitoXpress®-Intra O_2_-sensing probe [24] was from Agilent Technologies Ireland (Cork, Ireland). OptiMEM I medium, DAPI, TMRM, B27 serum-free supplement, TURBO DNAse, BCA^TM^ Protein Assay kit, RIPA lysis buffer, High-capacity cDNA reverse transcription kit, Fisher BioReagents™ EZ-Run™ Prestained Rec Protein Ladder and ProLong® Gold mounting medium were from Thermo Fisher Scientific (Waltham, MA). Amersham™ ECL™ Prime Western blotting reagent was from GE Healthcare Life Sciences (Waukesha, WI), pre-made acrylamide gels, running and transfer buffers, and 5× sample buffer were from GenScript (Piscataway, NJ). Complete protease inhibitor cocktail tablets and phosSTOP phosphatase inhibitor tablets were from Roche (Mannheim, Germany). RNA isolation RNeasy Plus Universal Mini Kit was from Qiagen (Venlo, Netherlands). SensiFAST™ SYBR® Lo-ROX Kit for qPCR was from Bioline (London, UK). DMOG, DCA, NAO, mitochondrial fission inhibitor Mdivi-1, antimycin A, myxothiazol, FCCP, TEMPOL, NAC, L-cysteine, SDT, McCoy’s 5A, RPMI, DMEM, DMEM-F12 Ham, collagen IV, PDL and all other reagents were from MilliporeSigma (Burlington, MA). Plasticware was from Sarstedt (Ireland), Corning Life Sciences (Corning, NY), Ibidi (Martinsried, Germany), and Greiner Bio One (Frickenhausen, Germany).

### Tissue Culture

Human colon cancer HCT116 wild type (WT) and SCO2-deficient cells [22] were kindly provided by P.M. Hwang (NIH, Bethesda, MD). Unless stated otherwise, cells were maintained for up to 14 days in McCoy’s medium supplemented with 10% FBS, 2 mM L-Glutamine, 100 U/ml penicillin / 100 μg/ml streptomycin (P/S), and 10 mM HEPES (pH 7.2) in humidified atmosphere of 5% CO_2_ / 95% air at 37°C under: 1) ‘normoxia’ (∼19% O_2_) in a regular CO_2_ incubator; 2) chronic hypoxia, ChH (3% O_2_ or <0.1% O_2_) in a hypoxia workstation (Coy Laboratory Products, Grass Lake, MI). Along with ChH, in the experiments with primary neurons CIH was used; neurons were grown at 3% O_2_ in a CO_2_/O_2_ incubator (Thermo Fisher Scientific) with passages conducted at the atmospheric O_2_ levels (20.9% O_2_, ∼30 min).

For all cell cultures, passages were performed every 3-4 days. During incubation cells were harvested for Western blotting analysis and other assays.

Treatments with compounds affecting respiration, free radical production, redox state and mitochondrial biogenesis were conducted as described in Results section. DMSO solution was used as Mock treatment when compounds were prepared using this solvent.

The density of HCT116 cell seeding for different protocols depended on their type and growing conditions and has been selected empirically for individual protocol. This was particularly challenging because seeding had to be done 2-3 days prior to the experiment. Thus, SCO2-deficient cells proliferated slower that WT cells, though having a larger size. Under ChH, proliferation rate of WT cells decreased, opposite to SCO2-deficient cells [25]. For protein and RNA isolation cells were seeded at 4×10^5^ (WT) and 5×10^5^ (SCO2-deficient) cells per well on 6-well plates. For mitochondrial protein isolation, RNA-seq, and Ribo-seq analyses, cells were seeded on 15 cm Petri dishes at 5-7×10^6^ (WT) and 7-8×10^6^ (SCO2-deficient) For immunostaining and live cell imaging, 1.2×10^4^ (WT) and 1.5×10^4^ (SCO2-deficient) cells per cm^2^ were seeded on glass-bottom dishes and inserts (Ibidi). In all cases cells were seeded on the surfaces pre-coated either with 0.01% collagen IV (plastic) or a mixture of 0.007% collagen / 0.003% PDL (glass) and then grown for 24-48 h. Collagen and PDL were prepared in weak acetic and boric acid solutions, respectively.

### Animal and primary cell models

All protocols involving animals were approved by the University College Cork Animal Experimentation Ethics Committee; experiments were conducted under licence from the Irish Government in accordance with national and EU legislation (European Directive 2010/63/EU).

#### 1. Animal model of chronic sustained hypoxia

Tissue samples were dissected from animals as reported in [26]. Briefly, age- and weight-matched adult male C576Bl/J mice (Charles River Laboratories, UK; n = 32) were exposed to six weeks of normoxia, chronic sustained hypoxia (10% O_2_, or FiO_2_ = 0.1), or chronic sustained hypoxia with antioxidant supplementation with TEMPOL or NAC (4 groups: n=8 per group) in environmental chambers (OxyCycler Model A84, BioSpherix Ltd, USA) with precise control of ambient O_2_ concentration. Mice were housed conventionally on a 12:12-h light–dark cycle at room temperature with food and water available *ad libitum*. Environmental chambers were opened briefly once a week for routine husbandry. At the end of the treatment periods, animals were anaesthetised by 5% isoflurane inhalation (in O_2_) and euthanised by cervical dislocation followed by immediate dissection of tissue samples. Brain cortex samples were snap-frozen and stored at -80°C. For Western blotting analysis tissues were lysed using RIPA buffer.

#### 2. Primary neuronal culture

Neuronal cell culture was prepared as described [27]. Briefly, cortices from E18 rat embryos (Biological Services Unit, UCC) were dissected; neural stem cells were dissociated and maintained in a monolayer on the bottom of PDL-coated 25 cm^2^ flasks (Corning) in DMEM/F12 Ham medium supplemented with L-Gln (2 mM), P/S (1%), B27 (2%), and 1% FBS. Cells were grown for 8 days at 19% and 3% O_2_ as above; medium was replaced regularly every 3 days. Proteins for Western blotting analysis were isolated using a standard RIPA lysis buffer.

### Generation of mRNA-seq libraries for RNA-seq and Ribo-seq analyses

Cells grown on 15 cm Petri dishes were quickly washed with PBS containing cycloheximide (100 μg/ml), scraped in polysome lysis buffer (20 mM Tris-HCl, pH 7.5, 250 mM NaCl, 1.5 mM MgCl_2_, 1 mM DTT, 0.5% Triton X100, 100 μg/ml cycloheximide and 20 U/ml TURBO DNAse), collected in 1.5 ml Eppendorf tubes and lysed on ice for 10 min. After lysate clarification (10 min, 16000 g, 4°C), supernatants containing polysomes were collected.

The libraries for mRNA sequencing analysis, including total and ribosome-protected mRNA sequences, were generated as in [28]. Libraries were sequenced on an Illumina HiSeq 2000 system at the Beijing Genomics Institute (China).

### Analysis of mitochondrial cytochromes using Raman spectroscopy

Raman spectra were recorded from the cytoplasm regions at the boundary with the nucleus, on Raman InVia microspectrometer (Renishaw, UK) as described [29], with modifications. Raman spectra induced by the 532 nm laser were mostly produced by vibrations of heme bonds in b- and c-type cytochromes. Other major peaks corresponded to vibrations of phenylalanine residues (Phe) in proteins and C-C bonds in lipids (**Table S1**).

Under these excitation conditions, Raman scattering of oxidized cytochromes was negligible, and Raman spectra consisted mainly of reduced cytochromes of complex III (c, c1, b_high_ and b_low_). After recording the initial spectra from the cell in 3 randomly selected ROI, all available cytochromes (including cytochrome b of complex II) were fully reduced (1 a.u.) by incubating cells with SDT (5 min), and spectra were collected again from the same ROI. These signals were used to evaluate relative levels of reduced cytochromes at given experimental conditions.

### Immunofluorescence and Confocal Microscopy

For immunofluorescence analysis HCT116 cells were grown at 19% and 3% O_2_ as above, first in the flasks for 9 days and then in Ibidi microscopy multi-well chambers for 2 days. Cells were washed with PBS and fixed with 4% PFA. Immunostaining was performed as described [30], using antibodies against Cyt*c* and COXIV (primary), fluorescent secondary antibodies (Table S2) and DAPI counterstaining. The intensity and localisation of the target proteins were analysed on an Olympus FV1000 confocal laser scanning microscope, equipped with 405 nm, 488 nm and 543 nm lasers. DAPI, Alexa Fluor 488 and Alexa Fluor 555 fluorescence signals were collected with a UPLSAPO 60x/1.35 oil immersion Super Apochromat objective in sequential laser mode (to avoid spectral overlap) in up to 10 focal planes (0.5 µm steps), using standard settings: DAPI - 405 nm / 420–460 nm, Alexa Fluor 488 – 488 nm / 500–540 nm, Alexa Fluor 555 – 543 nm / 560–620 nm (excitation / emission). The resulting single plane differential interference contrast (DIC) and z-stacked fluorescence images were processed using FV1000 Viewer software (Olympus) and Adobe Photoshop.

### Live Cell Confocal Microscopy

Loading of the cells with 20 nM TMRM (ΔΨm probe) and 2 μM NAO (CL dye, ΔΨm sensitive [31]) was performed in OptiMEM I medium for 30 min. TMRM was maintained in the medium at 20 nM during the measurement. FCCP prepared in OptiMEM was prewarmed prior to the addition to the samples. Imaging was performed on an Olympus FV1000 confocal laser scanning microscope under controlled CO_2_, humidity and temperature conditions, using standard excitation / emission spectra (488 nm / 510-540 nm and 543 nm / 550-600 for NAO and TMRM, respectively). Data collection and processing were performed as for immunofluorescence analysis.

### Protein Isolation and Western Blotting Analysis

Whole cell lysates were prepared in RIPA lysis buffer and analysed as described [32]. Briefly, clarified cell lysates with equal protein concentrations were separated using 4-20% polyacrylamide gels (GenScript, NJ), transferred onto a 0.45 μm Immobilon^TM^-P PVDF membrane (MilliporeSigma) using wet mini-transfer system Hoefer™ TE 22 (Hoefer, CA) and probed with antibodies (**Table S2**), which were prepared in 5% (w/v) BSA or non-fat dry milk in TBST. Immunoblots were analysed with: 1) HRP-conjugated secondary antibodies and ECL™ Prime reagents using the LAS-3000 imager (Fujifilm, Japan) or 2) fluorescent secondary antibodies using Odyssey® infrared imager (LI-COR Biosciences, Cambridge, UK). Levels of holo-Cyt*c* were analysed using its intrinsic peroxidase activity [33], ECL™ Prime reagents and the LAS-3000 Imager, or dedicated antibodies. Images were processed with ImageJ program.

### Isolation of Mitochondrial Proteins

Mitochondrial fraction was isolated from HCT116 cells according to [34]. Briefly, cells were washed in PBS containing 1 mM glucose (to ensure glycolytic ATP flux and steady mitochondrial membrane potential), collected, pelleted and stored at -80°C. Pellets were homogenised using isolation buffer (0.25 M sucrose, 10 mM Tris, pH 7.5, 1 mM EDTA) and an ice cold glass homogeniser (Wheaton® Glass 7 mL Dounce Tissue Grinder, Thermo Fisher Scientific). After removal of cell debris (centrifugation at 600 g, 4°C), crude mitochondrial fraction was obtained (centrifugation at 10,000 g, 4°C) and lysed using RIPA buffer. Proteins were prepared for Western blotting analysis as above.

### Isolation of Total RNA and RT-PCR Analysis

Total RNA was isolated using RNA isolation kit (Promega), as per manufacturer protocols. Reverse transcription reaction was performed using 2 µg of total mRNA and ImProm-II™ RT System (Promega, Madison, WI). SYBR Green based qPCR was conducted on the AB7300 machine and analysed using the 7300 System SDS Software (Applied Biosystems, CA); reaction was controlled for the absence of genomic DNA amplification. Each sample was analysed in triplicate. Primers were designed using Primer-BLAST program (http://www.ncbi.nlm.nih.gov/tools/primer-blast/) (**Table S3**).

### Flow cytometry analysis of apoptotic cells

HCT116 WT and SCO2-deficient cells grown on 6-well plates were treated for one hour with DCA, (100, 200, 300 and 400 μM). Cells were detached from the plates using accutase and counted. Then, 10^4^ cells were stained using Annexin V-FITC and propidium iodide (PI) for 20 min in the dark as per the manufacturer’s instructions (FITC Annexin V Apoptosis Detection Kit, BD Biosciences). Samples were analysed on a LSRII cytometer (Becton-Dickinson, Mountain View, CA). Fluorescence was detected using: 1) excitation at 488 nm and acquisition at 530/30 nm (FITC-Annexin V) and using excitation at 561 nm and acquisition at 610/20 nm (PI).

### Analysis of cellular O_2_ levels

WT and SCO2-deficient cells were maintained at different atmospheric O_2_ for 7 days, loaded with 10 μg/ml MitoXpress®-Intra probe [24], trypsinised, seeded at 0.5-1.5×10^4^ per well on 96-well plates pre-coated with collagen IV and grown for another 24 h under either 19% or 3% O_2_. iO_2_ analysis was performed using TR-F reader Victor 2 (PerkinElmer, MA), which was placed in a hypoxia chamber adjusted to 19% or 3% O_2_ at 37°C. Each sample well was measured repetitively every 3-5 min over 3 h, taking two intensity readings at delay times of 30 and 70 µs and gate time 100 µs (excitation / emission: 340/642 nm). These intensity signals were converted into phosphorescence lifetime and then to O_2_ levels.

### Statistical Analysis and Data Presentation

Statistical analysis was performed using the results of 3-4 independent experiments. Confidence levels of ≥ 0.01 were deemed as statistically significant. To ensure the accuracy and fidelity of the data, the experiments were performed in 3-8 replicates.

For Western blotting and PCR, results were normalised to α-tubulin and β-actin, respectively. Protein phosphorylation levels were normalised to the total content of the corresponding protein.

The differences between SCO2-deficient and WT cells (N, ChH, CIH) in TMRM, NAO, protein, mRNA and other measured parameters were evaluated using non-parametrical Mann-Whitney U-test. Unless otherwise stated, protein and mRNA levels were related to corresponding values in WT cells grown at 19% O_2_ (1 a.u.). Similarly, Cyt*c* protein levels were normalised for neuronal culture and brain cortex tissue samples. Graphically, the normalised results are presented as average values ± standard deviation (for kinetic experiments) or box-and-whisker plot. Boxes show medians and interquartile ranges (IQR), whiskers delineate the highest and lowest values in the samples, as long as these values are not outliers. Values higher than Q3+1.5xIQR or lower than Q1-1.5xIQR are considered outliers (filled circles).

## Results

### Cyt*c* protein levels is increased in COX-deficient and hypoxic cells

Previously, we reported that chronic hypoxia induces a substantial increase in Cyt*c* levels in neural PC12 and SH-SY5Y cells [35]. The concomitant decrease in respiration observed under hypoxic conditions suggested modulation of mitochondrial Cyt*c* content to facilitate electron transfer to COX. Based on this rationale, SCO2-deficient HCT116 cells, which lack electron flux from COX to O₂, were selected as a model of impaired electron transfer. Alone with SCO2-proficient (wild type, WT) SCO2-deficient cells were cultured continuously at 50–90% confluence either in a standard CO₂ incubator (∼19% atmospheric O₂, hereafter termed “normoxia”) or in a hypoxia workstation set to 3% or ∼0.1% O₂.

Under normoxic conditions, total Cyt*c* protein content was markedly elevated in SCO2-deficient cells compared with WT cells (**Fig. 1a**) and did not change further upon continuous culture at 3% O₂. In contrast, in SCO2-proficient HCT116 cells grown under identical conditions, Cyt*c* levels increased progressively in response to hypoxia, ultimately reaching levels comparable to those observed in SCO2-deficient cells (**Fig. 1b, c**). Exposure of either cell line to more severe hypoxia (∼0.1% O₂) did not result in any further increase in Cyt*c* levels (**Fig. 1d**). We therefore concluded that, in SCO2-deficient cells, Cyt*c* levels are already maximized under ‘normoxic’ conditions and cannot further respond to changes in O₂ availability. In WT cells, however, the actual intracellular O₂ levels could be substantially lower than the atmospheric 3% due to high O₂ consumption [36]. Using an O₂-sensitive probe MitoXpress®-Intra and a plate reader Victor2 placed inside a hypoxia chamber, we found that under static culture conditions at 3% atmospheric O₂, the O₂ concentration in SCO2-proficient cells decreased to ∼1% (8–11 µM), whereas in SCO2-deficient cultures O₂ levels remained close to atmospheric values (**Fig. S1**). Notably, when SCO2-proficient cells were returned from 3% O₂ to 19% O₂, Cyt*c* levels declined to their original normoxic values within 3-4 days (**Fig. 1e**).

**Figure 1.**
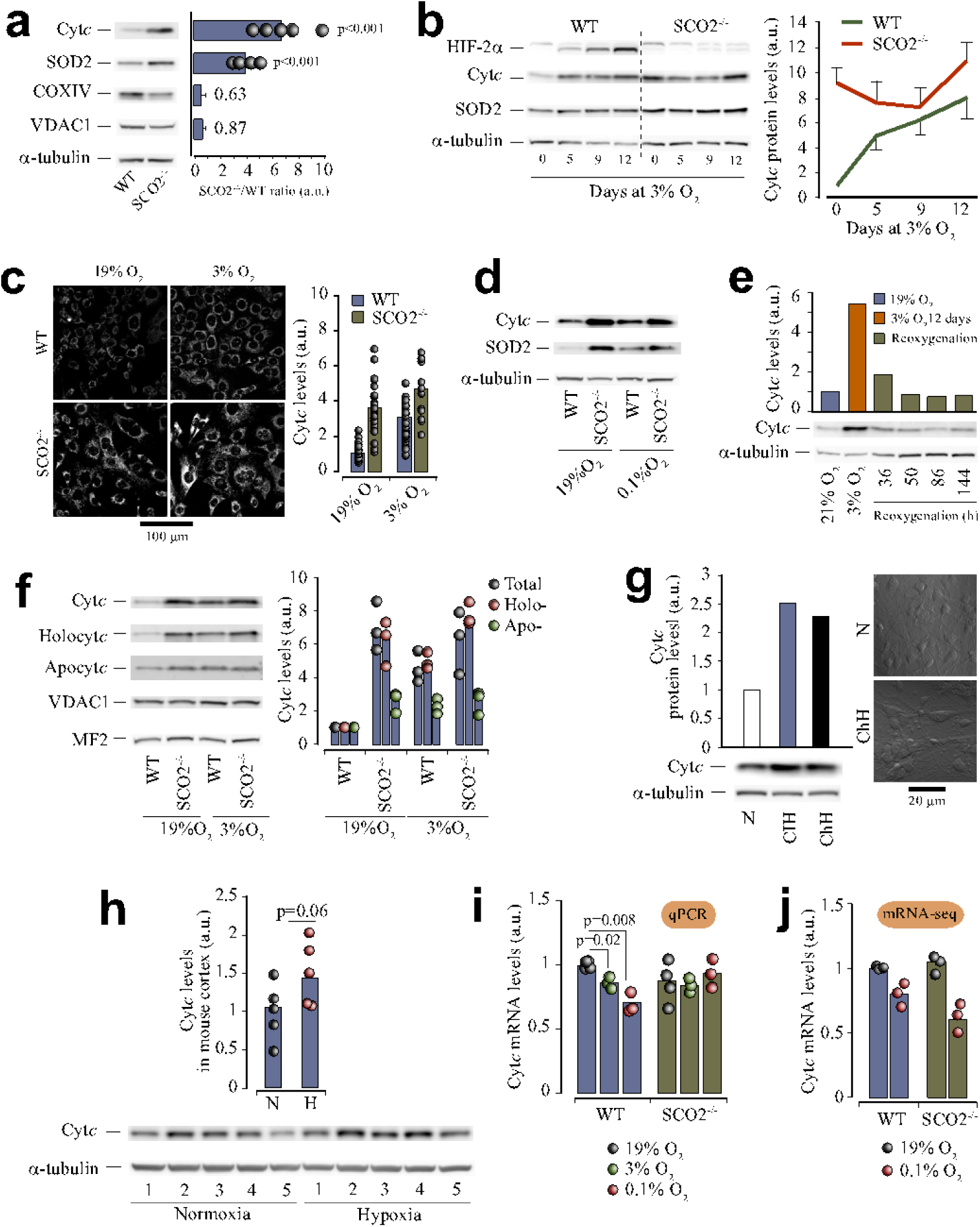
Cytochrome c protein levels increase in low respiring cells in a gene expression-independent manner. **a.** Western blotting analysis of mitochondrial protein levels in SCO2-deficient and SCO2-proficient HCT116 cells. Unlike COXIV, Cyt*c* and SOD2 levels increase, compared to the levels of a ‘housekeeping’ VDAC1 protein. **b**. Western blotting analysis of dynamics in Cyt*c* accumulation in WT and SCO2-deficient cells over 12 days of growing in hypoxic conditions for 12 days. Changes in HIF-2α confirm effects of chronic hypoxia (3% O_2_) on cells. **c**. Immunostaining analysis of Cyt*c* levels in WT and SCO2-deficient cells grown at 19% and 3% O_2_ for 9 days; five confocal planes taken with a 0.5 μm step are overlaid. **d**. Western blotting analysis of Cyt*c* and SOD2 levels in cells exposed to 19% and 0.1% O_2_ for 9 days. **e**. Effects of reoxygenation on Cyt*c* protein levels in SCO2-proficient HCT116 cells grown at 3% O_2_ for 12 days. **f**. Western blotting analysis of Cyt*c* sub-populations (total, holo- and apo-) in mitochondria of WT and SCO2-deficient cells grown at 19% and 3% O_2_ for 9 days. **g**. Changes in Cyt*c* protein levels in primary neuronal cell grown for 9 days in chronic and intermittent hypoxia (N=1). DIC images of the cells are shown. **h**. Analysis of Cyt*c* protein levels in cerebral cortex tissues of chronically hypoxic and normoxic mice (N=5 in each group). **i**. qPCR analysis of Cyt*c* mRNA levels in SCO2-deficient and SCO2-proficient HCT116 cells exposed to 19%, 3% and 0.1% O_2_ for 9 days (N=3 in each condition). **j**. mRNA-seq analysis of *CYSC* transcription levels in SCO2-deficient and SCO2-proficient HCT116 cells grown at 19% and 0.1% O_2_ for 9 days (N=1).

Immunostaining analysis of HCT116 cells revealed that changes in Cyt*c* abundance driven by hypoxia and SCO2 deficiency occurred predominantly in mitochondria (**Fig. 1c**). Consistently, Western blotting analysis of mitochondrial proteins demonstrated that it was primarily holo-Cyt*c* that accumulated in responses to SCO2 deletion and hypoxia (**Fig. 1f**). In SCO2-deficient cells, chronic hypoxia did not lead to any further increase in holo-Cyt*c* levels. Apo-Cyt*c* levels were also elevated in SCO2-deficient cells and, to a lesser extent, in hypoxic SCO2-proficient cells. To discriminate between apo- and holo-Cyt*c* on immunoblots, holo-Cyt*c* (a heme-conjugated protein) was visualised via its intrinsic peroxidase activity, whereas apo-Cyt*c* was detected using specific primary antibodies and fluorescent secondary antibodies (LI-COR IRDye 800CW).

The effect of chronic hypoxia on Cyt*c* abundance was further examined in actively respiring primary cell and tissue models. An upward trend in Cyt*c* levels was observed in cortical neurons from embryonic rats cultured for 8 days at 3% O₂ under either sustained or intermittent hypoxic conditions **(Fig. 1g, h**). Similarly, Cyt*c* content tended to increase in cerebral cortex tissue from mice exposed to sustained hypoxia for 6 weeks (**Fig. 1i**). As reported previously [26, 37], increased haematocrit and right ventricular mass confirmed a robust systemic response to chronic hypoxia in these animals.

### Analysis of potential contributors to Cyt*c* accumulation

Next, we examined several factors that could contribute to Cyt*c* accumulation by increasing either its production de novo or its cellular lifespan. These included: (a) enhanced Cyt*c* transcription and translation; (b) favourable redox state of Cyt*c*; (c) increased CL availability; (d) inhibition of electron flow through Cyt*c*; (e) altered ROS production; (f) activation of hypoxia-inducible factor (HIF) signalling; and (g) altered mitochondrial remodelling.

#### a. Transcription and translation of Cytc

Two independent mRNA-based approaches, qPCR and RNA-seq, revealed no increase in *CYCS* transcription in SCO2-deficient and hypoxic WT cells (**Fig. 1j**). Moreover, in SCO2-proficient cells, *CYCS* transcription was reduced upon continuous exposure to 0.1% O_2_. Consistently, ribosome profiling (ribo-seq analysis) did not reveal upregulation of Cyt*c* mRNA translation in SCO2-deficient or hypoxic SCO2-proficient HCT116 cells (**Fig. S2b**). Notably, a reduced polysome density was observed in SCO2-deficient cells, suggesting lower overall protein synthesis compared with SCO2-proficient cells (**Fig. S2a**).

#### a. Redox state of Cytc

Inhibition of electron transfer from Cyt*c* to complex IV is expected to increase the ferro-Cyt*c* / ferri-Cyt*c* ratio. Therefore, next we compared the redox state of Cyt*c* in normoxic SCO2-deficient and proficient HCT116 cells using Raman spectroscopy, which allows discrimination between reduced and oxidised forms of Cyt*c* [29]. Consistent with our immunoblotting data, Raman analysis confirmed elevated Cyt*c* levels in SCO2-deficient cells (**Fig. 2a, b**). In addition, spectral profiling revealed an increased ratio of total Cyt*c* to cytochrome b (Cytb), indicating a specific relative enrichment of Cyt*c*, as a representative of cytochrome family within the mitochondrial electron transport chain.

**Figure 2.**
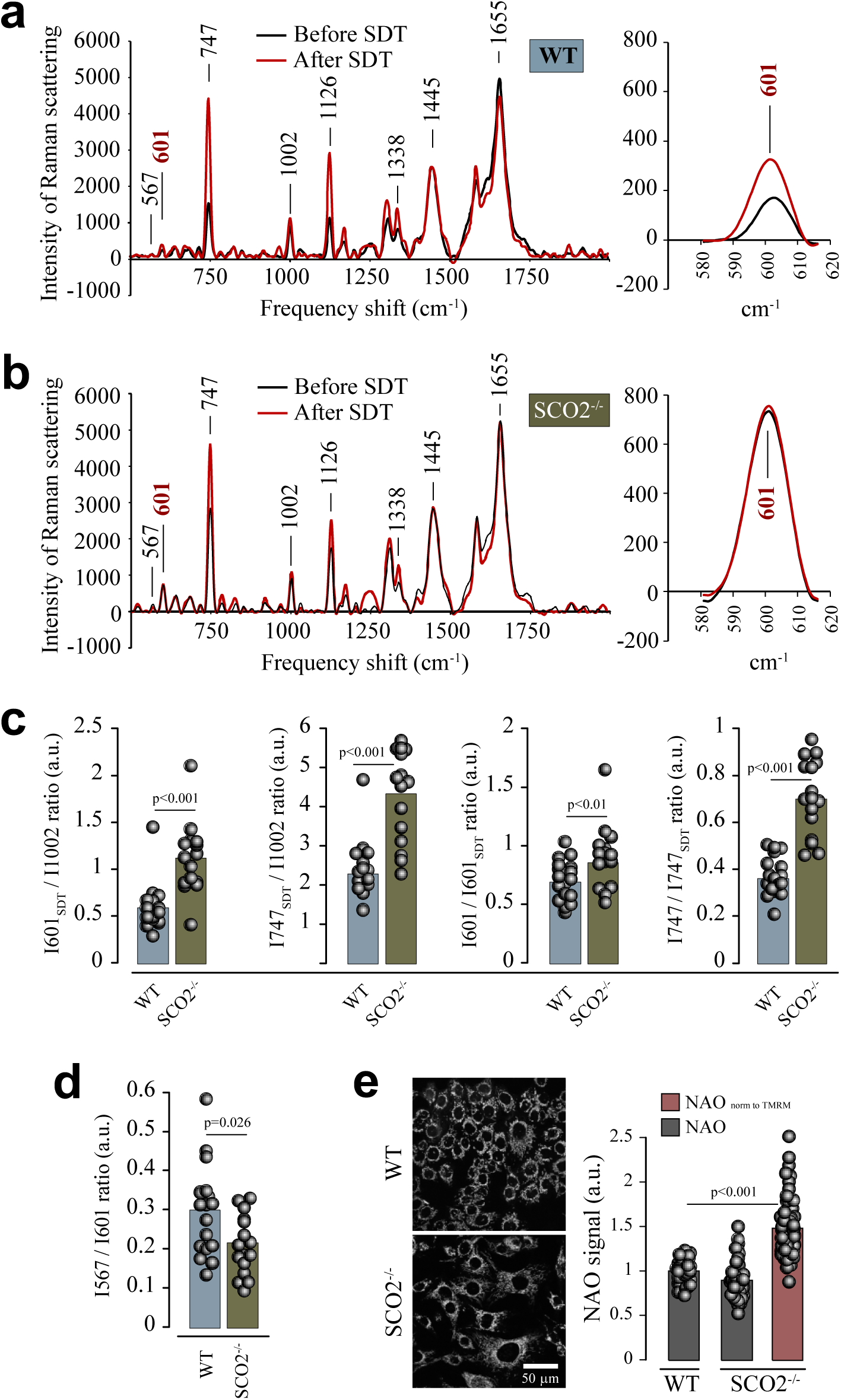
Analysis of Cyt*c* levels, redox state and interaction with the mitochondria in HCT116 cells. **a**, **b**. Raman spectra of WT and SCO2-deficient cells taken before and after addition of dithionite (SDT). **c**. Relative amounts of total cellular Cyt*c* vs total protein (shown as a ratio of I601/I1002 and I747/I1002) and reduced Cyt*c* vs total Cyt*c* (I601/I601_SDT_ and I747/I747_SDT_) **d**. The ratio of the levels of reduced Cyt*c* unbound to the mitochondrial membrane vs total reduced Cyt*c* (I567/I601). **e**. Levels of mitochondrial CL measured using a confocal live cell microscopy of NAO (2 μM, 30 min). In SCO2-deficient cells NAO fluorescence signals were also normalised to TMRM signals (not shown), which previously demonstrated that ΔΨm in SCO2-deficient cells is decreased by 40%; N = 18 (WT) and N = 16 (SCO2-deficient) cells.

We further found that the fraction of reduced Cyt*c* was significantly higher in SCO2-deficient cells compared to WT cells (**Fig. 2c**), consistent with impaired electron transfer to COX. This observation aligns with previous reports demonstrating increased structural and thermal stability of reduced Cyt*c* [38]. Moreover, the fraction of the membrane-unbound ferro-Cyt*c* was decreased in SCO2-deficient cells, suggesting enhanced retention of Cyt*c* at the inner mitochondrial membrane in non-respiring mitochondria (**Fig. 2d**).

#### b. Cardiolipin (CL) availability

The mitochondrial phospholipid CL, which is essential for maintaining mitochondrial structural and functional integrity, represents the primary interaction partner of Cyt*c* [39]. We therefore hypothesised that elevated CL levels might facilitate Cyt*c* accumulation by increasing both the probability and strength of Cyt*c*–membrane interactions [40]. To assess CL abundance, we employed NAO, a fluorescent dye reported to bind CL selectively [41]. Comparable NAO fluorescence signals were observed in WT and SCO2-deficient cells under resting conditions (**Fig. 2e**). However, NAO accumulation within mitochondria was strongly dependent on mitochondrial membrane potential (ΔΨm), because fluorescence rapidly dissipated upon mitochondrial depolarisation with the uncoupler FCCP (**Fig. S3**), in agreement with reported observations [31]. To account for differences in ΔΨm between cell lines, NAO signals in resting WT and SCO2-deficient cells were normalised to tetramethylrhodamine methyl ester (TMRM) fluorescence. As we demonstrated previously using a mitochondrial potential-dependent dye TMRM, SCO2-deficient cells exhibit an approximately 40% reduction in ΔΨm, compared to WT cells [42]. After ΔΨm normalisation, the data indicated a moderate increase in CL levels in cells lacking SCO2, which may enhance Cyt*c* accumulation.

#### c. Electron transport through Cytc

Both conditions associated with Cyt*c* accumulation—COX deficiency and severe hypoxia—are characterised by reduced electron flux through Cyt*c*. To assess whether any impairment in electron transfer, not necessarily associated with COX deficiency, is sufficient to induce Cyt*c* accumulation, we investigated the long-term effects of selective pharmacological inhibition of complex III in HCT116 cells using antimycin A and myxothiazol.

Phenotypically, cells treated continuously for 9 days developed features resembling SCO2-deficient HCT116 cells, including reduced proliferation rates and increased cell size, while exhibiting only minor changes in mitochondrial membrane potential (**Fig. S4**). Western blotting analysis revealed elevated Cyt*c* levels in SCO2-proficient cells following complex III inhibition, whereas no further increase in Cyt*c* abundance was observed in SCO2-deficient cells treated with antimycin A or myxothiazol (**Fig. 3a**). In contrast, treatment with the mitochondrial uncoupler FCCP, which enhances electron flux in functional mitochondria, did not alter cell morphology (**Fig. S4**) and Cyt*c* content (not shown) in SCO2-proficient cells.

**Figure 3.**
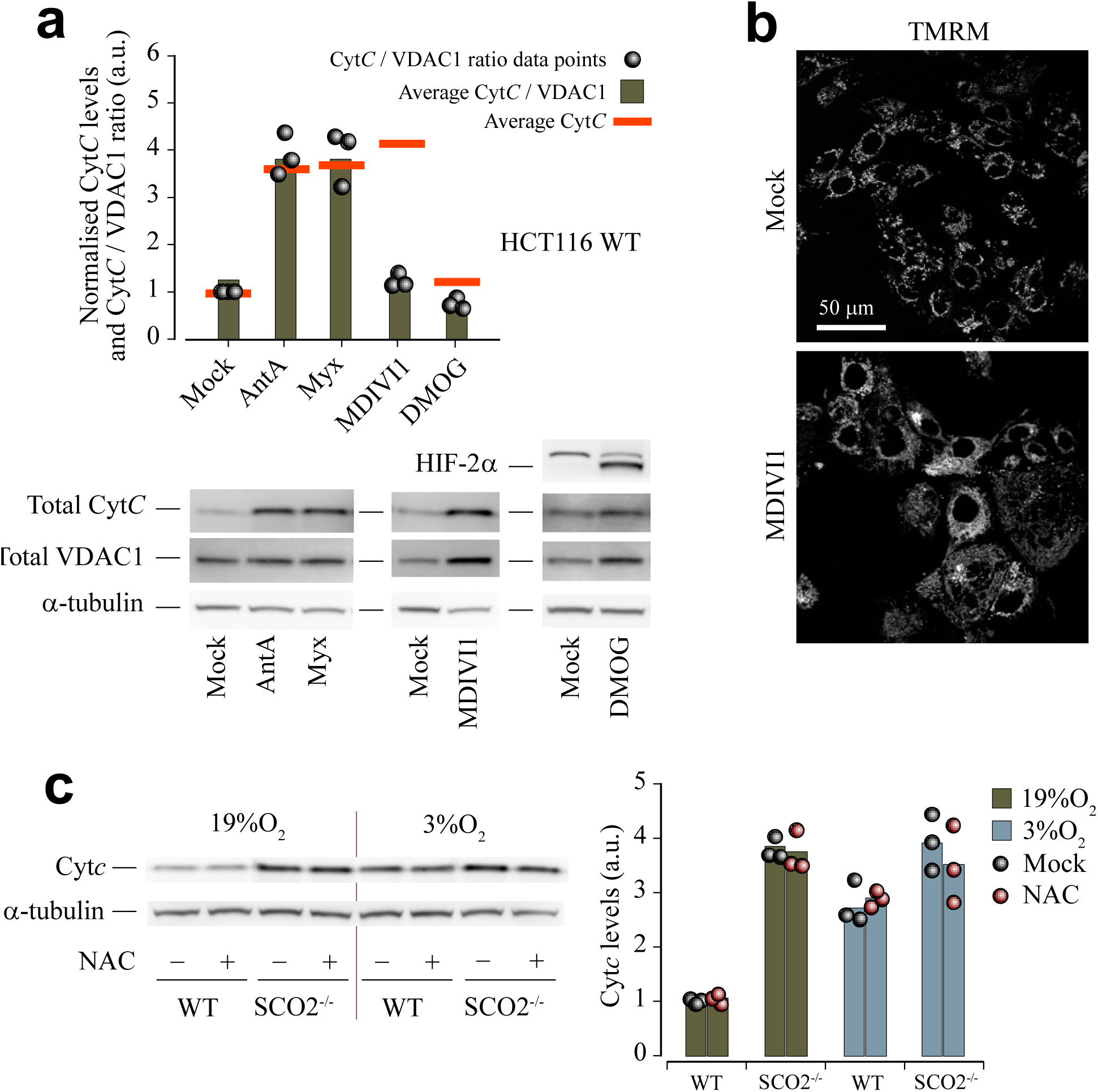
Effects of mitochondrial modulators on Cyt*c* levels in HCT116 cells. **a**. Western blotting analysis of the effects of mitochondrial complex III inhibitors antimycin A (Ant A, 5 μM) and myxothiazol (Myx, 2 μM), and mitochondrial remodelling factors Mdivi-1 (5 μM) and DMOG (1 mM), on Cyt*c* and VDAC1 protein levels, all drugs being applied for 9 days. HIF-2α levels show that DMOG treatment mimics effects of hypoxia on HIF-dependent processes in cells. **b**. Immunofluorescence analysis of the effects of Mdivi 1 on mitochondrial tubulation; live cell confocal imaging of TMRM staining. c. Western blotting analysis of the effects of NAC, an inhibitor of ROS, on Cyt*c* protein levels. N = 3 in (**a-c**).

Raman spectroscopy further confirmed an increased relative abundance of ferri-Cyt*c* in WT cells treated with antimycin A (**Fig. S5**), consistent with previous reports, showing elevated oxidation of Cyt*c* upon complex III inhibition [43].

#### d. Free radical overproduction and scavenging

It has been reported that SCO2-deficient cells overproduce reactive oxygen species (ROS), and that ROS generation decreases under low O₂ availability, coinciding with improved cell survival and proliferation (42). At the same time, the direction of changes in ROS production under hypoxia remains debated (e.g., [44, 45]). Given that Cyt*c* readily reacts with free radicals and participates in redox signalling, oxidative damage and detoxification, ROS-dependent mechanisms could, in principle, contribute to Cyt*c* accumulation in SCO2 deficient and hypoxic SCO2-proficient cells [25]. To test this possibility, we examined the effects of antioxidants on Cyt*c* levels in HCT116 cells and in vivo. Treatment with NAC, a widely used ROS scavenger, had no effect on Cyt*c* accumulation induced either by COX deficiency or by chronic hypoxia (3% O₂) in HCT116 cells (**Fig. 3c, d**). Likewise, neither NAC nor the superoxide dismutase mimetic Tempol altered the moderate increase in Cyt*c* content observed in the cerebral cortex of mice subjected to chronic hypoxia (**Fig. S6**). These findings are consistent with our observation that Ant A and Myx elicited similar effects on Cyt*c* levels (Fig. 3a). Given that these agents have opposing effects on mitochondrial ROS production, this provides additional evidence that ROS do not contribute to the observed effect (**Fig. 3a**).

#### e. Activation of HIF signalling

Cytochrome c levels are not known to be directly regulated by hypoxia-inducible factor (HIF) signalling. However, HIF activation could indirectly contribute to Cyt*c* accumulation by modulating the expression of factors involved in Cyt*c* turnover. We investigated this possibility using SCO2-proficient and SCO2-deficient cells chronically exposed to the α-ketoglutarate competitor DMOG, which suppresses HIF-α degradation and thereby activates HIF signalling in HCT116 cells [30].

Under normoxic conditions, SCO2-deficient cells contained elevated levels of HIF-α, consistent with metabolic inhibition of its degradation [46]. Chronic exposure to deep hypoxia further increased HIF-α abundance in these cells. In SCO2-proficient cells, both chronic hypoxia and continuous DMOG treatment stabilised HIF-1α and HIF-2α and induced the expression of canonical HIF target genes [30, 46]. In parallel, cell morphology and mitochondrial network organisation in hypoxic and DMOG-treated cells became similar to those observed in SCO2-deficient cells (**Fig. S7**), in agreement with previous reports demonstrating enhanced mitochondrial fusion downstream of HIF activation [47], However, despite robust activation of HIF signalling, DMOG treatment did not alter Cyt*c* protein levels in SCO2-proficient cells (**Fig. 3a**), in contrast to chronic hypoxia. These findings indicate that HIF activation alone is not sufficient to drive Cyt*c* accumulation.

#### f. Contribution of mitochondria remodelling

Although HIF-dependent mitochondrial fusion did not appear to contribute to Cyt*c* accumulation, we next examined whether altered mitochondrial dynamics could nonetheless influence Cyt*c* levels. Inhibition of mitochondrial fission with Mdivi-1 led to pronounced mitochondrial tubulation (**Fig. 3b**) and increased levels of VDAC1, a marker of mitochondrial mass (**Fig. 3a**). While total Cyt*c* levels were also elevated under these conditions, normalisation to VDAC1 revealed no change in Cyt*c* abundance relative to mitochondrial content (**Fig. 3a**). These data, alone with a negligeable effect on Cyt*c* levels of DMOG, an activator of fusion (**Fig. 3a**), suggest that mitochondrial remodelling is unlikely to be a primary driver of the Cyt*c* accumulation observed in SCO2-deficient and hypoxic cells.

Together, these results indicate that Cyt*c* accumulation in cells with compromised mitochondrial respiration is driven primarily by increased protein stability and lifespan, rather than by enhanced de novo synthesis. Cyt*c* levels increase independently of its redox state, ROS production, HIF signalling, gene expression levels or mitochondrial remodelling. In turn, reduced electron flux through Cyt*c* along with enrichment of the inner mitochondrial membrane with CL, which helps stronger / longer retention of Cyt*c* in mitochondria, likely act as important stabilising factors promoting Cyt*c* accumulation.

### SCO2-deficient cells are resilient to pro-apoptotic stimuli

We previously demonstrated that loss of SCO2 in HCT116 cells is associated with an increased dependence on AKT signalling and that pharmacological inhibition of AKT with MK2206 induces pronounced apoptosis in SCO2-deficient cells [46]. Given that elevated Cyt*c* levels could be associated with enhanced type II (Cyt*c*-mediated) apoptosis, we next examined apoptotic signalling in both HCT116 cells grown in CIH conditions.

Apoptosis was assessed by monitoring the degradation of PARP, a widely used marker of caspase activation. Under normoxic conditions, basal PARP levels were significantly lower in SCO2-deficient cells than in WT cells (**Fig. 4a, p** < 0.01, t-test), likely reflecting sustained PARP cleavage in response to chronic oxidative stress, consistent with previous report [48]. Exposure to CIH resulted in a progressive reduction of PARP levels in SCO2-proficient cells, whereas in SCO2-deficient cells CIH did not alter PARP degradation (**Fig. 4b**). Under ChH, PARP levels remained largely unchanged in both cell lines.

**Figure 4.**
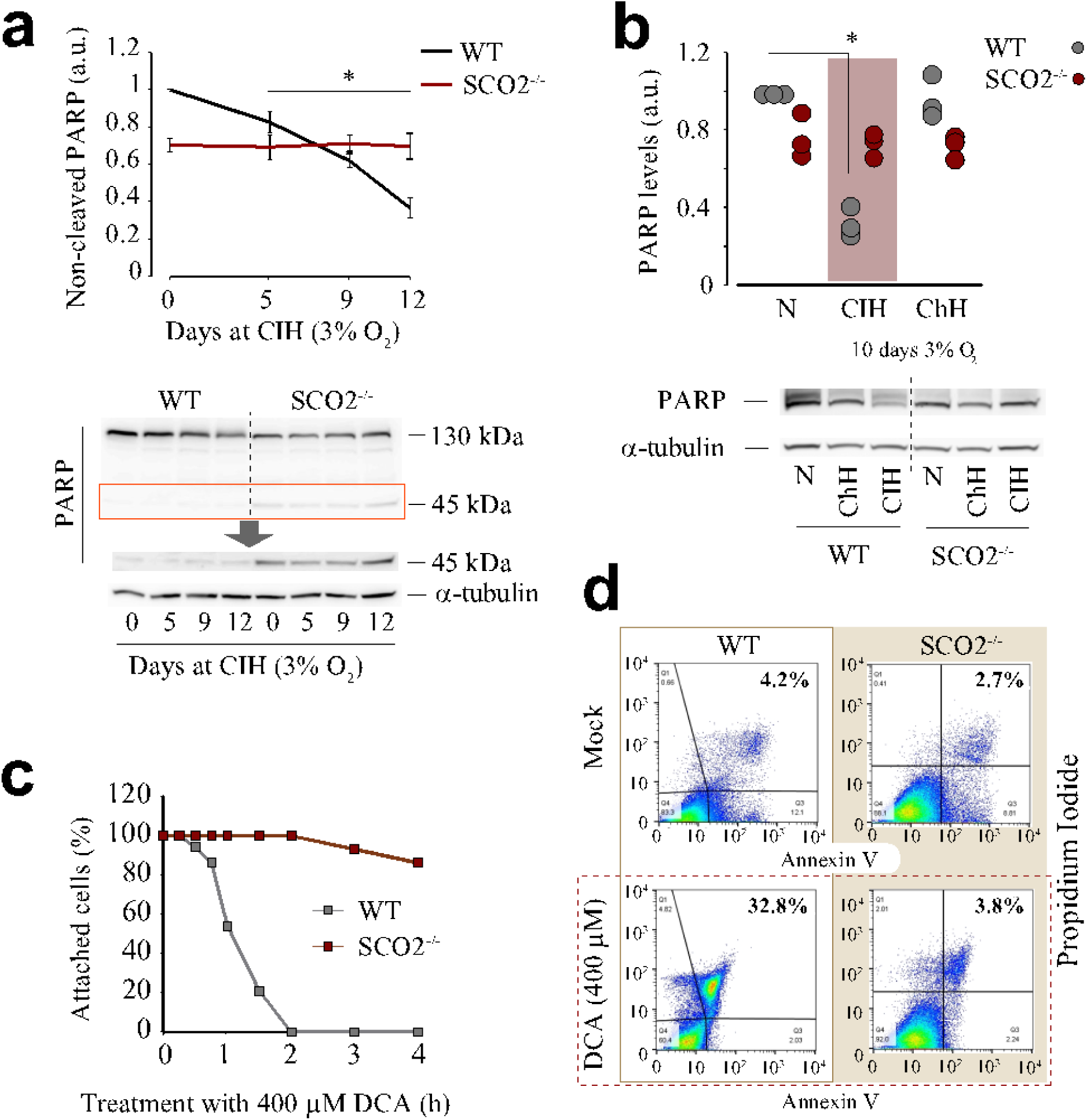
Increased sustainability of SCO2-deficient cells to pro-apoptotic stimuli. **a.** Dynamics of PARP levels in WT and SCO2-deficient cells chronically exposed to intermittent hypoxia (CIH, 10 days). A 45 kDa band, which corresponds to a product of PARP degradation (highlighted by green rectangle in the main blot), is shown in the bottom panel using a longer exposure time. **b.** Different effects of CIH and ChH on PARP degradation; changes in PARP levels caused by CIH are highlighted in the quantitative plot. **c.** Time-dependent effect of deoxycholic acid (DCA, 400 μM) on viability of HCT116 cells. **d.** FACS analysis of the levels of cell death induced in HCT116 cells by 400 mM DCA (1 h treatment). In a, b and d, N = 3; asterisks show significant difference between hypoxic and ‘normoxic’ WT cells (t-test): in (A), p = 0.003, p = 0.0005, and p < 0.0005 for 5, 9 and 12 d hypoxia; in (B), p = 0.00006; in (C), N = 4, p = 0.02 (15 min), p = 0.004 (30 min), p = 0.0007 (1 h), and p < 0.0005 (>1 h).

To further assess the resistance of SCO2-deficient cells to pro-apoptotic stimuli, we treated both cell lines with increasing concentrations of DCA, a potent inducer of Cyt*c*-dependent apoptosis in HCT116 cells [49]. Quantification of cell attachment to collagen-coated culture plates revealed a striking contrast between the two cell types. SCO2-proficient cells exhibited a concentration- and time-dependent loss of adhesion following DCA treatment, whereas SCO2-deficient cells remained largely attached over a wide range of DCA concentrations and treatment durations (**Fig. S8** and **Fig. 4c**, respectively). Consistently, flow cytometric analysis using Annexin V and propidium iodide demonstrated a marked reduction in viability of SCO2-proficient cells, but not SCO2-deficient cells, following exposure to 0.4 mM DCA (**Fig. 4d**).

Collectively, these results indicate that, despite elevated mitochondrial Cyt*c* levels, SCO2-deficient cells exhibit reduced susceptibility to Cyt*c*-mediated apoptosis in response to common pro-apoptotic stimuli, including intermittent hypoxia and DCA.

## Discussion

In this study, we demonstrate that Cyt*c* protein levels increase in cells and tissues with compromised mitochondrial electron transfer, including COX-deficient cells and cells exposed to chronic hypoxia. Importantly, this accumulation is not driven by increased *CYCS* transcription or translation (Fig. 1i, j and Fig. S2) but instead reflects enhanced protein stability and mitochondrial retention. Our findings reveal a previously underappreciated adaptive response of mitochondrial Cyt*c* to impaired electron flux, uncoupled from the ‘classical’ apoptotic signalling.

In principle, mitochondrial import efficiency and heme availability could influence steady-state Cyt*c* levels; however, the directionality of these effects argues strongly against enhanced Cyt*c* biosynthesis as an explanation for the accumulation observed in SCO2-deficient and hypoxic cells. Apocytochrome c is translocated into mitochondria via the TOM (translocase of the outer mitochondrial membrane complex and subsequently stabilised through covalent heme attachment by holocytochrome c synthase (HCCS) in the intermembrane space. Importantly, efficient Cyt*c* import and maturation are favoured by intact mitochondrial membrane potential, balanced redox conditions, and adequate heme supply [50, 51].

Under conditions of impaired electron transport, including SCO2 deficiency and chronic hypoxia, mitochondrial membrane potential is partially reduced, and mitochondrial redox homeostasis is altered [52]. Both factors are expected to impair, rather than enhance, TOM-dependent protein translocation and heme utilisation. Consistent with this, mitochondrial dysfunction has been widely associated with reduced protein import capacity and compromised maturation of intermembrane space proteins [53]. Likewise, heme synthesis and trafficking are tightly coupled to mitochondrial metabolic state, and disruptions in respiratory activity have generally been linked to restricted heme availability rather than increased heme flux [54].

Taken together with the absence of *CYCS* transcriptional or translational upregulation, these considerations strongly argue against increased Cyt*c* production or improved import as drivers of Cyt*c* accumulation in our models. Instead, they strengthen the conclusion that Cyt*c* accumulates despite cellular conditions that would normally limit its biogenesis. This apparent paradox further supports a model in which reduced electron flux and enhanced membrane association prolong Cyt*c* lifespan by suppressing its turnover, thereby outweighing any concomitant constraints on Cyt*c* synthesis or mitochondrial import.

Our observation that SCO2 deficiency and hypoxia both lead to elevated Cyt*c* levels supports the idea that reduced electron flux through the ETC stabilises Cyt*c*. Raman spectroscopy revealed an increased proportion of reduced (ferro-) Cyt*c* in SCO2-deficient cells, consistent with impaired electron transfer to COX. Biophysical studies have shown that reduced cytochrome c adopts a more compact solution structure with a smaller radius of gyration compared with its oxidised counterpart, consistent with a relatively more stable conformation in the reduced state [55–57]. Moreover, analyses of redox-coupled dynamics indicate that the oxidised protein exhibits greater local unfolding and conformational flexibility than reduced Cyt*c*, which also implies differential stability between the two redox forms [58]. Thermodynamic measurements further show that the heme–methionine bond is more stable in ferrous cyt c than in its oxidised form, providing a mechanistic basis for enhanced stability of reduced Cyt*c* [56]. In addition, the decreased fraction of membrane-unbound Cyt*c* observed in SCO2-deficient cells suggests enhanced retention at the inner mitochondrial membrane, further limiting Cyt*c* turnover.

The principal mitochondrial phospholipid, negatively charged CL is responsible for anchoring Cyt*c* to the inner mitochondrial membrane and regulating its functional state. CL-Cyt*c* interactions are integral to both respiratory efficiency and apoptotic signalling: cytochrome c binds tightly to CL and can act as a CL oxygenase in apoptosis, and biophysical studies indicate conformational changes upon CL binding that stabilise membrane association [40, 59]. Although ΔΨm-corrected measurements indicated only a moderate increase in CL levels in SCO2-deficient cells, even subtle changes in CL abundance or distribution may substantially influence Cyt*c*–membrane interactions, supporting a model in which enhanced Cyt*c*–CL association contributes to mitochondrial Cyt*c* stabilisation when electron transfer capacity is limited.

In addition to CL-dependent mechanisms, reduction of ΔΨm may influence Cyt*c* behaviour under conditions of impaired respiration. Cyt*c* is a positively charged protein within the intermembrane space, whereas the inner mitochondrial membrane exhibits a strong electrical gradient generated by proton pumping during OxPhos. A reduction in ΔΨm in SCO2-deficient cells [60] could, in principle, alter the electrostatic environment at the intermembrane-space face of the inner membrane, potentially affecting Cyt*c*–membrane interactions. However, to date there is no direct experimental evidence demonstrating that loss of ΔΨm alone strengthens Cyt*c* binding to the inner mitochondrial membrane. Available studies primarily link ΔΨm dissipation to Cyt*c* release during apoptosis, a process that is mediated predominantly by CL oxidation and membrane permeabilisation rather than by electrostatic effects of membrane potential per se. Thus, while ΔΨm loss accompanies many physiological states in which Cyt*c* stabilisation is observed, its contribution is likely indirect and subordinate to other mechanisms. Pharmacological inhibition of complex III in SCO2-proficient cells with antimycin A or myxothiazol reduces electron flux upstream of Cyt*c*. Under these conditions, in the presence of atmospheric O₂, Cyt*c* becomes predominantly oxidised by COX, as confirmed by Raman spectroscopy (Fig. S…). Notably, despite this shift towards ferri-Cyt*c*, total Cyt*c* levels were also increased. These findings indicate that although Cyt*c* reduction favours its stabilisation, it is not the only, nor an absolutely necessary, condition for Cyt*c* accumulation. Rather, a sustained decrease in electron flow through Cyt*c*—whether due to inhibition upstream (complex III) or downstream (COX deficiency or hypoxia)—appears to slow Cyt*c* turnover and degradation, thereby promoting its accumulation.

Despite the well-established link between hypoxia and HIF signalling, our data indicate that activation of HIF alone is insufficient to induce Cyt*c* accumulation. Pharmacological stabilisation of HIF-α with DMOG recapitulated several characteristic features of the hypoxic response, including pronounced mitochondrial remodelling and fusion, in line with previous reports linking HIF signalling to altered mitochondrial dynamics [61, 62]. However, despite robust HIF activation and extensive mitochondrial fusion, DMOG treatment did not alter Cyt*c* protein levels. This dissociation indicates that Cyt*c* accumulation is not a direct consequence of HIF-dependent transcriptional reprogramming or HIF-driven changes in mitochondrial morphology.

To further dissect the contribution of mitochondrial dynamics per se, we independently manipulated mitochondrial morphology by inhibiting fission with Mdivi-1. Consistent with its established effects, Mdivi-1 induced marked mitochondrial elongation and increased mitochondrial mass, as reflected by elevated VDAC1 levels. Under these conditions, total Cyt*c* abundance was increased; however, normalisation to mitochondrial mass revealed no change in Cyt*c* content relative to the size of mitochondria. These findings indicate that mitochondrial remodelling and increased mitochondrial volume can passively elevate total Cyt*c* levels, but do not account for the selective accumulation of Cyt*c* observed in SCO2-deficient or hypoxic cells.

Our DMOG- and Mdivi-1-based experiments demonstrate that neither HIF-driven mitochondrial fusion nor altered fission–fusion balance is sufficient to drive Cyt*c* accumulation. Instead, these data support a model in which Cyt*c* levels are regulated primarily by mitochondrial bioenergetic state and electron flux through the respiratory chain, rather than by mitochondrial morphology or HIF-dependent transcriptional programmes.

Oxidative stress caused by ROS overproduction can influence conformation, interactions, localisation and activities of Cyt*c*, including its ability to induce apoptosis [63, 64]. However, neither antioxidant treatment nor pharmacological modulation of mitochondrial ROS production altered Cyt*c* levels in our models. These findings align with recent work highlighting the context-dependent nature of mitochondrial ROS signalling in hypoxia and respiratory chain dysfunction [65, 66]. Collectively, our data argue against ROS as a major determinant of Cyt*c* accumulation under conditions of impaired electron transport.

Strikingly, despite elevated mitochondrial Cyt*c* levels, SCO2-deficient cells displayed reduced susceptibility to Cyt*c*-mediated apoptosis in response to intermittent hypoxia and DCA. This observation challenges the simplistic view that increased Cyt*c* abundance necessarily enhances apoptotic competence. Although Cyt*c* release into the cytosol is a hallmark of type II apoptosis, increasing evidence indicates that Cyt*c* function is highly context-dependent and tightly regulated. Apoptosis requires not only Cyt*c* availability but also its posttranslational modifications and redox tuning, CL oxidation, mitochondrial outer membrane permeabilisation, and apoptosome assembly [6, 67, 68]. Our data suggest that, in respiration-deficient mitochondria of HCT116 cells, Cyt*c* is preferentially retained in a membrane-bound, reduced, and most likely, non-apoptogenic state, thereby uncoupling Cyt*c* accumulation from apoptotic execution.

It is important to note, though, that this conclusion is relevant to our cell model and experimental conditions, and the link between reduced respiration, Cyt*c* state and apoptosis cannot be generalised. However, there is evidence that other metabolic deficiencies, such as lack of fumarate hydratase activity, protect cells from pro-apoptotic stimuli and contribute to tumour progression in hereditary leiomyomatosis and renal cell cancer (HLRCC), where FH-deficient cells display resistance to apoptosis [69]. Similarly, deficiency or loss of succinate dehydrogenase (SDH) — another key mitochondrial enzyme linking the TCA cycle and ETC — is a characteristic of various SDH-deficient tumours and is associated with tumour progression, altered metabolic states, and poor clinical prognosis, suggesting that impaired respiratory complex II activity can support cancer cell survival and therapy resistance rather than enhance cytochrome c-dependent apoptosis [70].

In summary, we propose that Cyt*c* accumulation represents an adaptive mitochondrial response to impaired electron transfer, driven by reduced electron flux, enhanced membrane retention, and increased protein stability rather than increased synthesis. This mechanism preserves redox capacity while limiting apoptotic sensitivity, highlighting a protective reprogramming of Cyt*c* function in metabolically compromised mitochondria. These findings refine our understanding of Cyt*c* biology beyond its canonical roles and may have implications for hypoxia-associated pathologies, mitochondrial diseases, and tumour metabolism.

## Acknowledgement

Authors would like to thank Professor Ken O’Halloran (Head of Physiology Department, UCC) for supporting experiments on hypoxic animal tissue, and Dr. Anmol Khan (UCC) for technical support in RNA-seq and Ribo-seq data analysis. This work was supported by Science Foundation Ireland under Grants No. SFI/12/RC/2276_P2, 07/IN.1/B1804 to D.B.P., and Research Ireland Frontiers for the Future award (20/FFP-A/8929) to P.V.B.

## Abbreviations

Ant A: antimycin A
ChH: chronic hypoxia
CIH: chronic intermittent hypoxia
CL: cardiolipin
COX: cytochrome c oxidase, a.k.a. mitochondrial complex IV
Cyt*c*: cytochrome c
ΔΨm: mitochondrial membrane potential
DAPI: 4′,6-Diamidino-2-phenylindol
DCA: deoxycholic acid
DIC: differential interference contrast
DMOG: dimethyloxalylglycine
DMSO: dimethyl sulfoxide
ETC: electron transport chain
HIF: hypoxia inducible factor
NAC: N-acetyl cysteine
NAO: 10 N-nonyl acridine orange
OxPhos: oxidative phosphorylation
PARP: poly (ADP-ribose) polymerase
PDL: poly-D-lysine
SCO2: cytochrome c oxidase assembly protein 2
SOD2: superoxide dismutase 2 (mitochondrial)
SDT: sodium dithionite
TMRM: tetramethylrhodamine, methyl ester
VDAC1: voltage-dependent anion-selective channel protein 1

