## Supplemental Materials for "Mitochondrial cytochrome c accumulation accompanies reduced electron flux through complex IV without enhancing cell sensitivity to apoptosis"

Table S1: Assignment of main peaks in Raman spectra

| **Position, $cm^{-1}$** | **Description** | **Comments** |
|---|---|---|
| 567 | Out-of-plane deformation of heme in Cyt*c* | Is seen only in the spectra of Cyt*c* molecules that are not attached to the mitochondrial membrane |
| 601 | Bond vibrations in heme of c-type cytochromes | Characteristic and highly specific for *c*-type cytochromes in reduced state |
| 747 | Bond vibrations in heme of type *b* and *c* cytochromes | Main contribution from *c*-type cytochromes in reduced state |
| 1002 | Bond vibrations in phenylalanine | Can be used to estimate relative amount of protein and to normalise intensities of all Raman peaks * |
| 1126 | Bond vibrations in heme of type *b* and *c* cytochromes | Main contribution from *b*-type cytochromes in reduced state |
| 1338 | Bond vibrations in heme of type *b* and *c* cytochromes in reduced state | Characteristic and highly specific for *b*-type cytochromes |

* Relative amount of phenylalanine residues in proteins should not differ between samples

Table S2. List of antibodies

| **Company** | **Target / Host** | **Cat N** |
|---|---|---|
| Abcam, UK | Cytochrome c / Mouse | ab13573 |
| | Cytochrome c oxidase subunit IV (COX IV) / Rabbit | ab202554 |
| | Mitofusin 2 /Mouse | ab56889 |
| | Superoxide dismutase 2 (mitochondrial), Rabbit | ab13533 |
| R&D Systems, MN | Hypoxia inducible factor 2α (HIF-2α) / Goat | AF2886 |
| Santa-Cruz Biotechnologies, CA | Voltage-dependent anion-selective channel protein 1 (VDAC1) / Goat | sc-8828 |
| Sigma, MO | α-tubulin / Mouse | T5168 |
| | Anti-goat/sheep IgG-peroxidase / Mouse | A9452 |
| | Anti-rabbit IgG, peroxidase conjugated / Mouse | A1949 |
| | Anti-mouse IgG, peroxidase conjugated Goat | A0168 |
| Thermo Fisher Scientific, MA | Alexa Fluor 555 anti-mouse IgG (H+L) / Goat | A21424 |

Table S3. List of primers

| **Gene** | **Primer sequence** | **Product length (b.p.)** |
|---|---|---|
| CYSC | F: 5'-GGCAAGCACAAGACTGGGCCAA<br>R: 5'-TGCCCTTTCTTCCTTCTTCTTAATGCCG | 208 |
| ACTB | F: 5'-CGGCTACAGCTTCACCACCACG<br>R: 5'-AGGCTGGAAGAGTGCCTCAGGG | 205 |

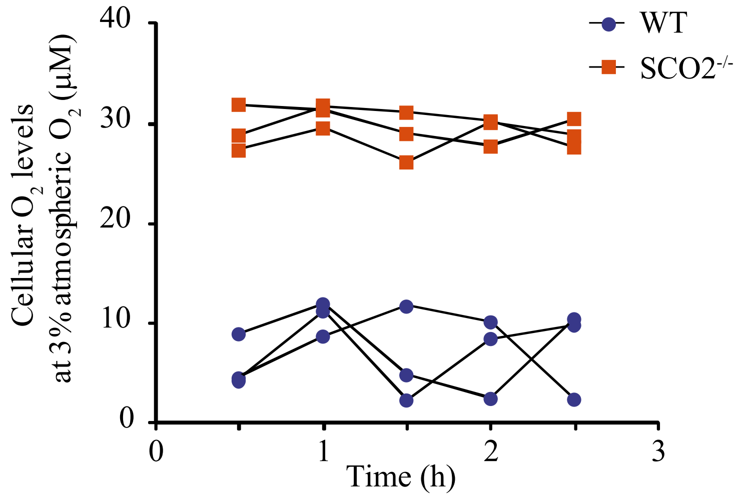


**Figure S1**. $O_2$ levels in HCT116 cells grown chronically under 3% atmospheric $O_2$. Measurements were taken every 30 min using MitoXpress®-Intra probe on a Victor2 plate reader placed in a hypoxia chamber. Results of 3 independent experiment are shown.

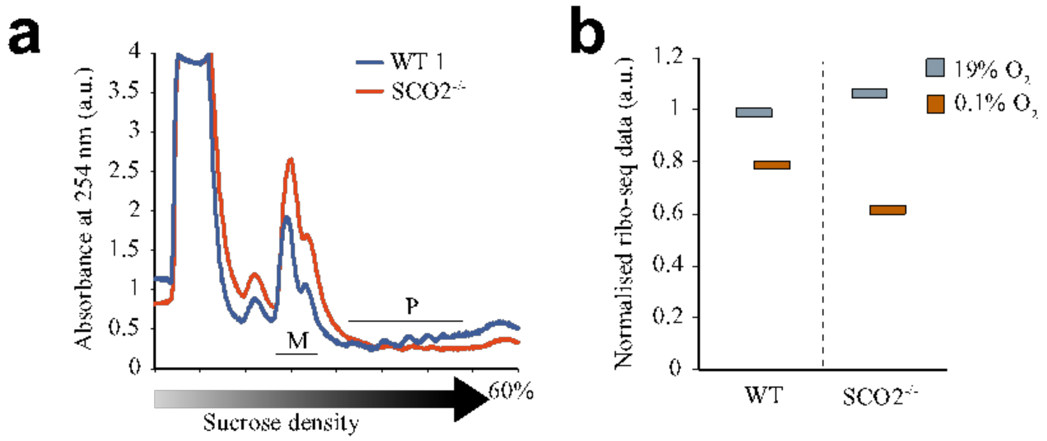


**Figure S2**. Ribosome density on mRNA extracted from HCT116 cells (a) and (b) normalised translation levels in SCO2-proficient and -deficient cells cultured for 9 days under normoxia (19% $O_2$) and deep hypoxia (0.1% $O_2$). In (a), ribosomal nucleoprotein complexes were separated by ultracentrifugation on a Beckman centrifuge in sucrose density gradient and detected by absorbance at 254 nm. 'M' and 'P' stand for monosome and polysome fractions, respectively. An increase in the M/P ratio indicates overall lower translation activity in the cells. In (b), results of ribo-seq analysis were normalised to the levels on Cyt*c* mRNA in normoxic WT cells; (N = 1).

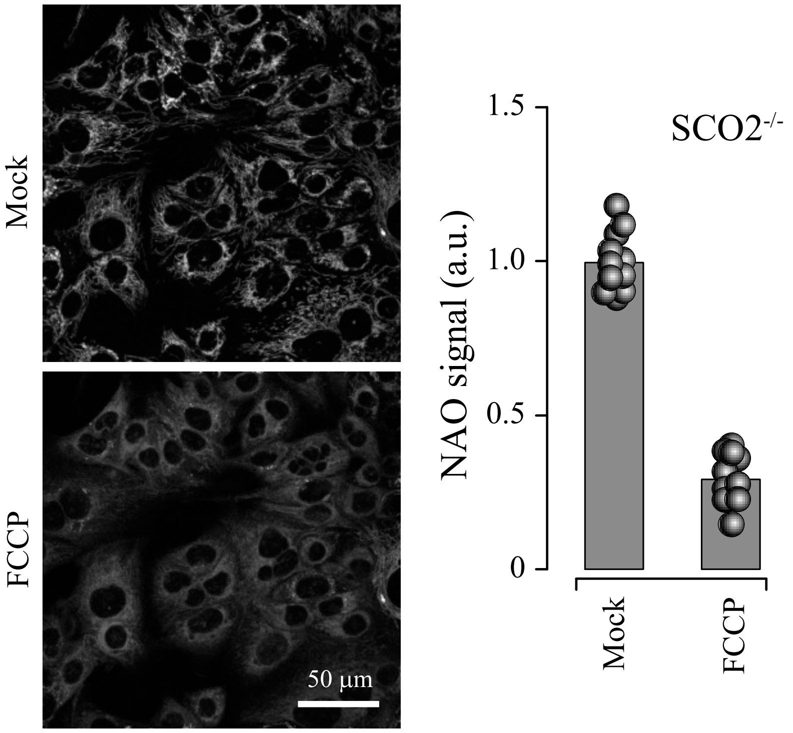


**Figure S3.** Changes in NAO fluorescence levels in HCT116 SCO2-deficient cell upon mitochondrial depolarisation (2 µM FCCP, 5 min). Images are stacks of four focal planes taken with a 0.5 µm step. N = 30.

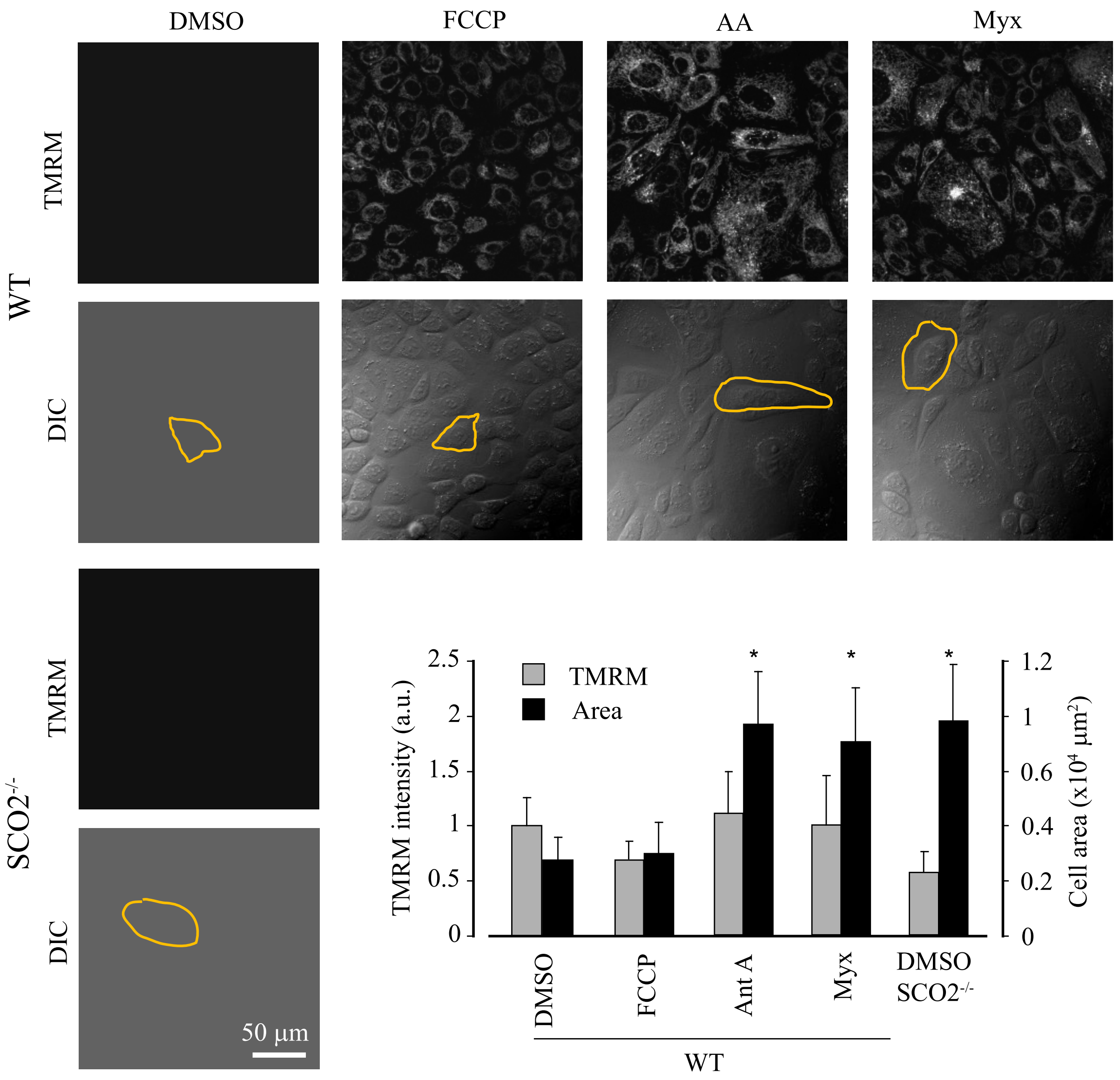


**Figure S4.** Effects of chronic FCCP, Ant A (AA) and Myx treatment on morphology and mitochondrial membrane potential of HCT116 cells. Live cell confocal imaging of TMRM fluorescence and cell morphology (DIC) of SCO2-proficient cells treated under normoxia for 9 days, compared to SCO2-deficient cells. Asterisks indicate significant difference from non-treated (mock) SCO2-proficient cells. Typical for each treatment cells are outlined.

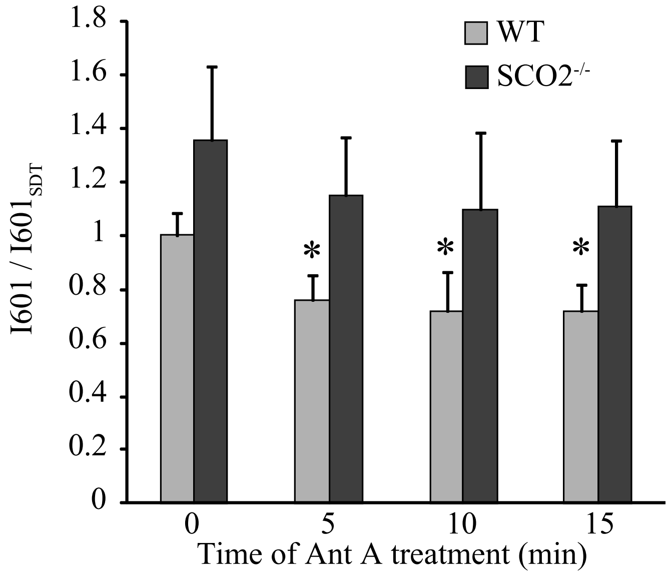


**Figure S5.** Effect of Ant A treatment on the ratio between reduced and oxidised Cyt*c* in HCT116 cells. Raman spectroscopy analysis of Cyt*c* oxidation state upon treatment with Ant A. $I601 / I601_{SDT}$ shows relative amount of reduced c-type cytochromes vs. total amount of c-type cytochromes. N = 3. Asterisks indicate significant difference from non-treated (mock) SCO2-proficient cells.

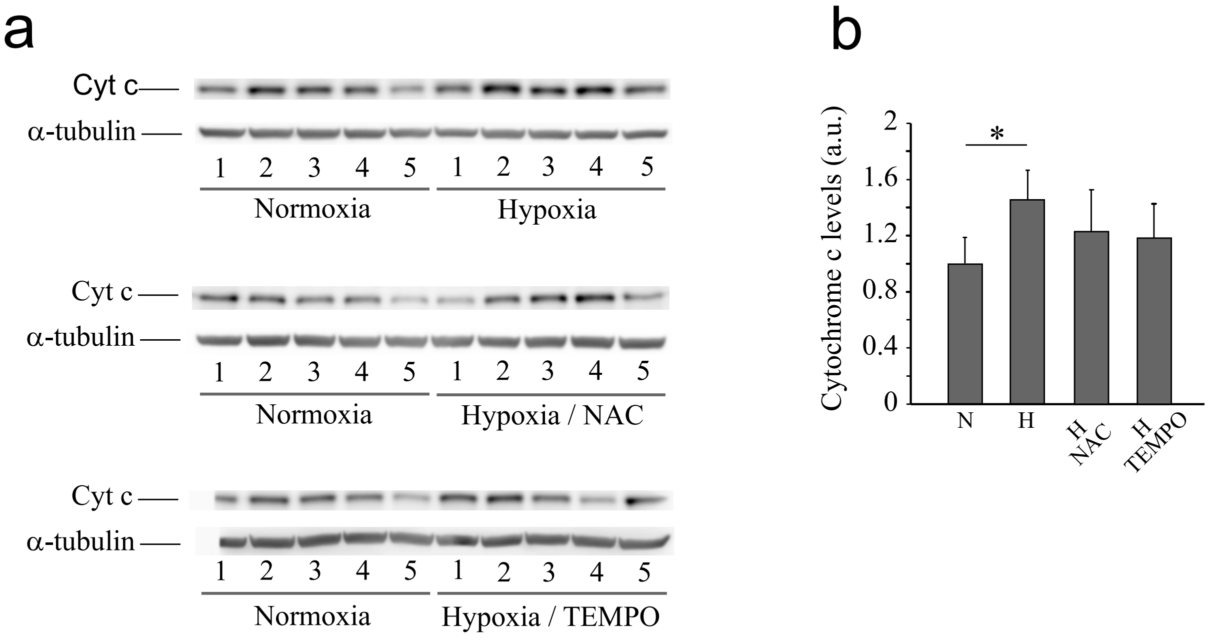


**Figure S6**. Effect of free radicals on accumulation of Cyt*c* protein in HCT116 cells and mouse cerebral cortex. A. Cyt*c* levels in HCT116 WT and SCO2$^{-}$deficient cells incubated for 10 days at 19% and 3% $O_2$ with or without 2 mM NAC. B. Cyt*c* levels in cortical tissue of mice, chronically subjected to reduced $O_2$ levels (10%, 6 weeks) with or without administration of Tempol or NAC in drinking water. N = 5 in each group. Asterisk shows significant difference.

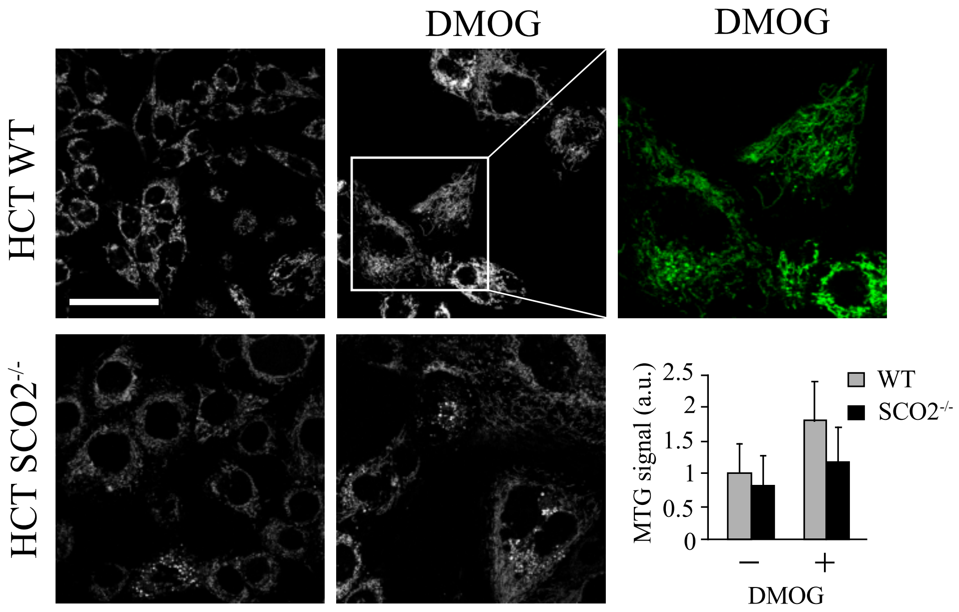


Figure S7. Changes in mitochondrial dynamics in HCT116 cell upon DMOG treatment for 9 days. Images of MTG fluorescence are stacks of four focal planes taken with a 0.5 μm step. Inset shows a typical 'tubular' structure of mitochondria in the treated WT cells. N = 28.

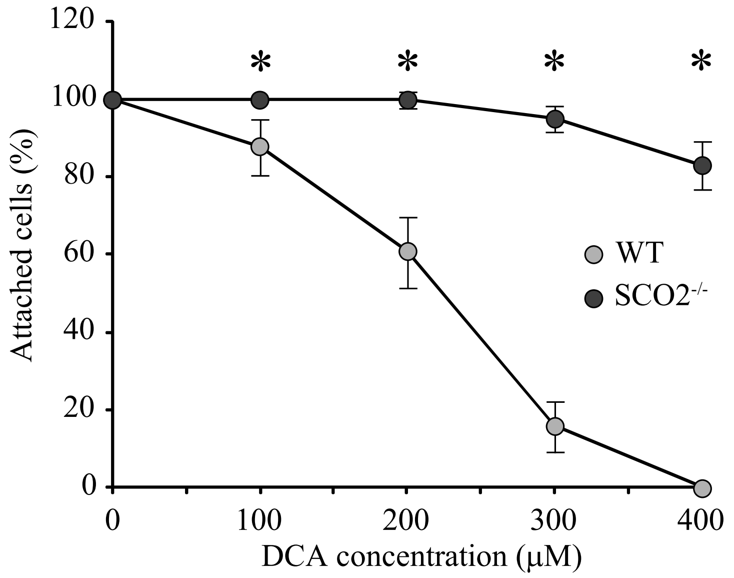


**Figure S8**. Concentration-dependent effect of DCA on viability of HCT116 cells; representative plot. Analysis was performed by counting cells attached to the surface of plastic after 2-h treatment. Asterisks show significant difference between DCA-treated SCO2-deficient and SCO2-proficient cells.
